# Small, fat-filled lipid droplets remain spherical as they indent a nucleus, dilute the lamina, and cause rupture

**DOI:** 10.1101/2022.09.01.506190

**Authors:** Irena L. Ivanovska, Michael P. Tobin, Lawrence J. Dooling, Dennis E. Discher

**Author notes:** equal contribution (Irena L. Ivanovska, Michael P. Tobin).

## Abstract

The nucleus in many cell types is a stiff organelle, and yet fat-filled lipid droplets (FD’s) in the cytoplasm can be seen to indent and displace the nucleus. FD’s are phase-separated liquids with a poorly understood interfacial tension γ that determines how FD’s interact with other organelles. Here, micron-sized FD’s remain spherical as they indent both the nucleus and peri-nuclear actomyosin, dilute Lamin-B1 locally independent of Lamin-A,C, and trigger rupture with locally persistent accumulation in the nucleus of cGAS, a cytosolic DNA sensor. FD-nucleus interactions initiate rapid mis-localization of the essential DNA repair factor KU80, and nuclear rupture associates with DNA damage and perturbed cell cycle. Similar results are evident in FD-laden cells after constricted 3D-migration, which is impeded by FD’s. Spherical shapes of small FD’s are consistent with a high γ that we measure for FD’s mechanically isolated from fresh adipose tissue as ∼40_mN/m – which is far higher than other liquid condensates, but typical of oils in water and sufficiently rigid to disrupt cell structures.

## Introduction

Many cell types have fat-filled lipid droplets (FD’s), and in some such as fat tissue cells the FD’s abut the nucleus (Farese and Walther, 2009). Fat is also soft to the touch, but its high levels of Lamin-A,C that confers nuclear stiffness and strength are surprisingly close to levels in rigid bone rather than the low levels in soft marrow or brain (Swift et al., 2013). High Lamin-A,C in fat thus hints at a curious rigidity of FD’s, especially given that Lamin-B1 is relatively invariant across tissues including fat. Deeper insight might eventually help clarify how fat is lost with aging of some stiff tissues (e.g. skin), and with lipodystrophy caused by some drugs (i.e. HIV protease inhibitors (Anuurad et al., 2010)) and mutations that suppress mature lamin-A (Bidault et al., 2011).

The nucleus is reportedly stiffer than the micron-size FD’s in mesenchymal stem cells (MSCs) undergoing adipogenesis – based on the challenging approach of poking an FD in a cell with an AFM tip and then assuming FD’s are solid (Shoham et al., 2014). However, FD’s in MSCs visibly indent the nucleus (**Fig.1A-i**), and FRAP measurements have also suggested FD’s are fluid (e.g. (Lyu et al., 2021)). FD’s are often described as having an interfacial tension γ, or energy-per-area (units of mN/m), set by their oily triacylglycerols and sterol esters composition (Murphy and Vance, 1999),(Hsieh et al., 2012) plus a surrounding monolayer of ER-derived phospholipid and protein (Farese and Walther, 2009) (Thiam et al., 2013). The particular value of γ is important because the pressure (force-per-area) that an FD sustains and exerts against a nucleus, for example, is set by γ multiplied by the FD’s curvature (radius^−1^) (**Fig.1A-ii**); furthermore, *any distortion from a spherical shape* necessarily increases an FD’s surface area and costs an energy set by γ. Measurements of γ are currently known only for FD’s from sonicated lysates of kidney cells (Cos7 and HEK) and fly hemocyte-like embryonic cells, with γ = 2-5 mN/m for ∼3-10 µm diam FD’s (Ben M’barek et al., 2017), (Lyu et al., 2021) (**Fig.1Aii, scale**). For comparison, protein condensates have far lower γ ∼0.01 mN/m) and can be highly elongated in cells to “*conform to the local structure*” such as the cytoskeleton (Boddeker et al., 2022). A typical cell membrane will also quickly rupture under an effective tension of ∼10 mN/m (Evans et al., 1976), which implies that pressing a soft condensate and perhaps an FD against a cell membrane is unlikely to rupture a membrane.

**Figure 1.**
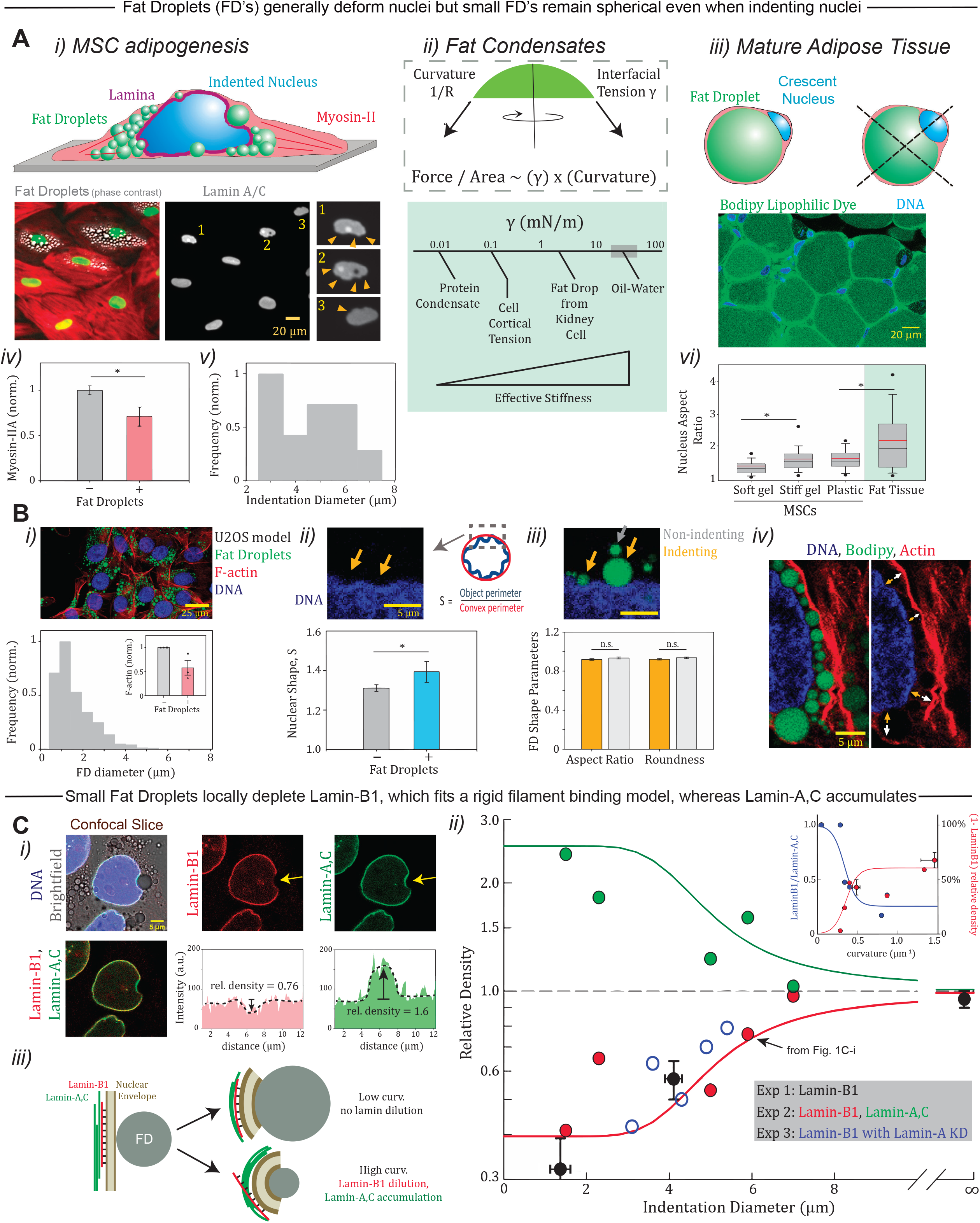
Fat droplets of varying curvature are sufficiently rigid to deform the nucleus and disrupt the nuclear lamina at sites of high Gaussian curvature. **A. i)** Adherent cell in culture filled with fat droplets (FD’s) portrays FD-induced indentation of the nucleus, as seen in fixed and immunostained MSCs undergoing adipogenesis (arrowheads). **ii)** Interfacial tension y multiplied by Gaussian curvature determines the pressure (force per area) that deforms an FD and nucleus; large FD’s deform at lower stress. A value of y was recently reported for FD’s isolated from one type of mammalian cell (Ben M’barek et al., 2017), which suggests a high value relative to other cell-scale tensions, such as the actomyosin mitotic cortex (Quang et al., 2021). Hexanol is partially miscible in water and γ = 6 mN/m, whereas long-chain hydrocarbon oils are insoluble with γ = 10-50 mN/m (Israelachvili, 2011). **iii)** Cells in fat tissue contain large FD’s that squeeze the nucleus to the cell periphery and deform it. **iv)** Actomyosin is suppressed in MSCs by FD’s (n>30 cells; p<0.05). **v)** Nuclear indentation diameter caused by FD’s varies within the ∼µm range (n>20 cells). **vi)** Nuclei in fat tissue have significantly higher nuclear deformation, exceeding even MSCs on stiff substrates (black=mean, red=median; n>250 cells; p <0.05). **B. i)** FD’s in U2OS cells after Oleic Acid addition (day-3) again show FD’s in the ∼µm range (1.5 μm med. diam.; N=2 exp; n>1000 cells) and suppression of F-actin (N=3 exp; n>30 cells). **ii-iv)** Nuclear indentations caused by FD’s show higher deviation from a smooth, convex shape (n>30 cells; p<0.05). However, both nuclear-indenting FD’s and distant, non-indenting FD’s maintain the same spherical shape (n>75 cells). FD’s that indent the nucleus appear back-stopped by actomyosin fibers, which FD’s also deform. **C. i)** Confocal mid-sections display nuclear indentation by small FD’s, and show lamin-B1 is locally depleted, while lamin-A,C is enriched. Immunofluorescence (IF) intensities are measured along the envelope. **ii)** Relative densities of lamin-B1 measured at sites of Gaussian curvature imposed by FD indentation fit a rigid filament binding model (see Methods; n=26, 5, 5 cells for the 3 expts). Lamin-A knockdown has no effect on Lamin-B1 results, but Lamin-A,C relative density fits the inverse of the same model. *Inset*: As functions of curvature, the probability of Lamin-B1 dilution also fits the model and Lamin B-to-A ratio fits well to the square of the model (see Methods). **iii)** Lamin-B1 and Lamin-A,C have distinct mechanical responses to indentation curvature: Lamin-B1 filaments resist bending and dissociate from the nuclear envelope at sites of high Gaussian curvature, whereas Lamin-A,C filaments accumulate.

Further consistent with nuclear indentation by an FD is the crescent shape of a nucleus at the periphery of mature adipocytes with a single, large FD filling the cytoplasm (**Fig.1A-iii**). Electron microscopy images convey a similar impression of indentation (Verstraeten et al., 2011). Whether FD’s are sufficiently rigid to indent a nucleus and perturb key functions such as DNA replication remains unclear but important because – even for nonmigratory settings of relevance to fat cells – nuclear rupture in interphase (De Vos et al., 2011), (Tamiello et al., 2013), (Nmezi et al., 2019) associates somehow with high nuclear curvature and with DNA damage (Cho et al., 2019), (Xia et al., 2018),(Pfeifer et al., 2018). In particular, Gaussian curvature (as the *product* of the two membrane curvatures) has been speculated to be of greater importance than Mean curvature (with a membrane flat in one direction), but nuclear rupture mechanics is complex, with failure likely driven in various ways. Nuclear indentation by FD’s provides a *physiologically relevant* means to examine possible consequences. Key results here show that small FD’s are sufficiently rigid to remain spherical when they indent and perturb the lamina’s density, favoring nuclear rupture, mislocalization of DNA repair factors, and increased DNA damage in both 2D culture and FD-impeded 3D migration.

## Results and Discussion

### FD’s displace and deform the cytoskeleton & nucleus, with small FD’s proving especially rigid

Adipogenesis of MSCs *in vitro* over ∼2 wks is seen to suppress actomyosin levels (**Fig.1A-iv**), consistent with known remodeling of the actin cytoskeleton (Cristancho and Lazar, 2011); (Spiegelman and Farmer, 1982), (Verstraeten et al., 2011). Actomyosin stress can drive nuclear deformation and even drive rupture when lamins are repressed (Tamiello et al., 2013), (Cho et al., 2019), (Hatch et al., 2013), (Xia et al., 2018), but nuclear indentation diameters that vary here from ∼3 to 7 μm clearly associate more directly with FD indentations than with actomyosin (**Fig.1A-v**). In mature fat tissue, the crescent-shaped nucleus is also highly elongated (**Fig.1A-vi**) consistent with a high local rigidity based on comparison with MSC nuclear elongation that increases with microenvironment stiffness (Boddeker et al., 2022), (Swift et al., 2013). These initial observations suggest an interplay of FD’s with the nucleus and cytoskeleton in tissue-relevant cells, but further insight could benefit from an experimentally tractable model system.

U2OS osteosarcoma cells are an MSC-derived lineage that have been used to study adipogenic processes (Mohseny et al., 2011), and they express more Lamin A,C than Lamin-B (Pfeifer et al., 2022). Adding high concentrations of monounsaturated Oleic Acid to U2OS cultures produces some FD’s in just a day (Brasaemle and Wolins, 2017) (Mannik et al., 2014); and FD’s of ∼1.5 μm median diameter eventually fill a similar area of cytoplasm as the nucleus (**Fig.1B-i**) and modestly suppress F-actin levels (**Fig.1B-i-inset**). Nuclear indentation in U2OS cells is again evident and readily quantified as a perimeter roughening (**Fig.1B-ii**).

FD’s that indent a nucleus are equally round whether indenting or not (**Fig.1B-iii**). Confocal z-stacks also show no significant flattening of FD’s, with profiles similar to spherical, polystyrene microbeads. Small FD’s are thus spherical (unlike large FD’s per **Fig.1A-iii**) and remain so even when they also induce a bend in actomyosin stress fibers adjacent to the nucleus. The notably thick stress fibers in U2OS cells (Hotulainen and Lappalainen, 2006) indeed bend (**Fig.1B-iv**), with effective diameters (<2 μm) that are small compared to the ∼10 μm persistence length of even a single actin filament (Gittes et al., 1993). Rescue of the indentations and distortions via cytoskeletal disruption is complicated by effects on cell morphology and a tendency to increase adipogenesis and FD’s (Cristancho and Lazar, 2011).

Confocal sections through FD’s that indent the nucleus reveal a major distinction between lamin intensities relative to adjacent undeformed regions: Lamin-B1 dilutes locally at indentation sites, whereas Lamin–A,C enriches (**Fig.1C-i**). Furthermore, Lamin-B1’s relative density decreases with indentation diameter across experiments, including with Lamin-A knockdown which reveals an independent response (**Fig.1C-ii**). Lamin filaments have a persistence length in situ that ranges from ∼0.2-1.5 μm (Turgay et al., 2017), and such rigidity could in principle frustrate attachment to the inner nuclear membrane of Lamin-B1 via its farnesylation groups that are lacking in mature lamin-A,C (Dechat et al., 2008). Such a single filament model of curvature-driven membrane detachment (see *Theory* in Methods) fits well to the Lamin-B1 measurements. Key to the fit is the curvature-dependent decrease in filament association energy (**Fig.1C-iii**), but an asymptote at high curvature (∼1.5 μm radius) suggests a limit to curvature-induced detachment-dilution and could relate to additional Lamin-B1 interactions. Experiments with GFP-Lamin-B Receptor (LBR) (not shown) that binds Lamin-B1 (Olins et al., 2010) again show FD-induced depletion, which adds confidence to the Lamin-B1 results; the LBR-demarked ER also does not accumulate at sites of FD indentation, which argues against a role for nucleus-linked ER in the local response. Lamin–A,C *enrichment* at sites of FD indentation (**Fig.1C-i**) seems consistent with Lamin-A,C lacking both LBR interactions and the lipid-driven association with membrane that is imparted to Lamin-B1 by its farnesylation.

Lamin–A,C’s density versus diameter of nuclear indentation is mathematically inverted from lamin-B1 dilution, so that the theoretical model is easily modified (see *Theory* in Methods) to fit well to the curvature-dependence of the ratio of lamin densities (**Fig.1C-ii inset**). Although Lamin-A,C’s response could be somehow compensatory for depletion of Lamin-B1 (**Fig.1C-iii**), U2OS cells with global knockdown of Lamin-B1 or Lamin-A,C show more frequent spontaneous nuclear rupture in standard 2D culture (Hatch et al., 2013),(Xia et al., 2018). Lamin-B1 knockout neurons likewise show increased nuclear rupture in vitro and in vivo (Chen et al., 2019), and lamin-A,C is nearly absent in neurons (Swift et al., 2013). Thus, given the abundance of and protection by lamin-A,C in U2OS cells, it was unclear whether the *localized* depletion of lamin-B1 caused by FD indentation increases nuclear rupture.

### Small Fat Droplets contacting the nucleus cause nuclear rupture, with loss of DNA repair factors

To begin to look for rupture, live cell imaging was done on FD-containing U2OS cells transfected with mCherry-cGAS (i.e. cyclic GMP-AMP synthase), which is a ‘DNA sensor’ that diffuses in the cytoplasm but will enter an interphase nucleus at sites of nuclear rupture and persistently bind chromatin for many hours (Harding et al., 2017), (Denais et al., 2016), (Raab et al., 2016). Live imaging over 3 h of cGAS-transfected cells with small FD’s showed nuclear foci of cGAS develop at sites of FD-nucleus interaction (**Fig.2A-i,ii**), and also showed such foci were comparatively rare in cells where FD’s are not near the nucleus (e.g. **Fig.2A-i**). Studies were done at day-1 with sparse FD’s to better relate a specific FD with a specific rupture site. The intensity of cGAS foci saturated by 20-40 mins, remained stable for >90 mins, and extended several µm’s from the site of FD-nucleus interaction (**Fig.2A-ii,iii**). Our observations seem consistent with mobile cGAS entering the nucleus and undergoing a liquid-to-solid gelation beyond ∼1 h (Du and Chen, 2018). Preexisting cGAS foci thus indicate past rupture events and remained stable over ∼hr’s, consistent with binding longevity (∼6-9 h), in ∼20% of FD-laden cells, which exceeds both the ∼10% observed during the 3 h of live-cell imaging and ∼5% seen as preexisting foci in cells with no FD’s near the nucleus (**Fig.2A-iv**). Multiple foci or ‘cGAS scars’ in a single nucleus of fixed cells with FD’s thus provide a ‘history’. Importantly, all cGAS entry events in live imaging were associated with FD’s < 3 µm, consistent with a relation to Lamin-B1 dilution (**Fig.1C**). Rescue of rupture-induced DNA damage and cell cycle delay is indeed achieved in part by overexpressing KU80, KU70 plus one more repair factor.

**Figure 2.**
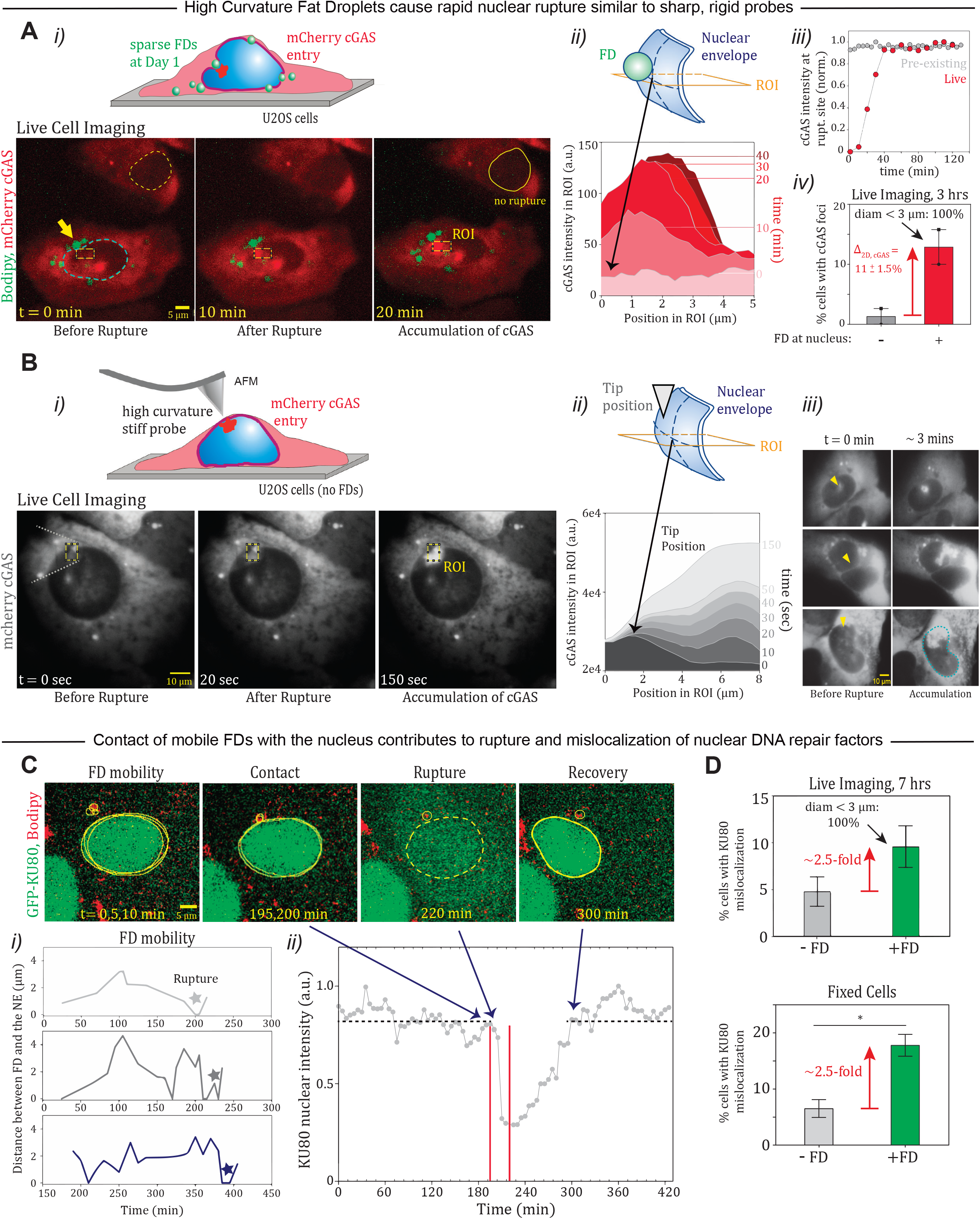
High curvature FD’s rupture the nucleus similar to stiff AFM probes, causing rapid accumulation of cGAS at rupture sites and mislocalization of DNA repair factor KU80 to cytoplasm. **A. i)** Cytosolic DNA sensor cGAS accumulates at sites of nuclear envelope rupture caused by high curvature FD’s, based on live cell experiments over 3 h (n=9 cells; dashed lines = nuclei). Cells were imaged 1-day after FD-induction. **ii)** Region of interest (ROI) intensity analysis at the rupture site (yellow rectangle) shows spatiotemporal accumulation of cGAS and reveals rapid accumulation by 10 min with saturation at 40 min. **iii)** cGAS intensity at rupture sites does not dilute for hours. **iv)** >10% of cells with FD’s at the nucleus show rupture during live imaging whereas <2% of cells rupture when FD’s are not nearby or present (n>30 cells per duplicate experiment). **B. i**,**ii)** A stiff AFM probe with high curvature pressed into contact also causes nuclear rupture and cGAS accumulation on μm scales over minutes. The plasma membrane does not rupture based on a constant intensity of cytoplasmic cGAS. **iii)** cGAS accumulates reproducibly within ∼3 mins of nuclear rupture. **C**. Mislocalization of DNA repair factor GFP-KU80 during live cell imaging (for 7 h) occurs frequently upon contact with moving FD’s (n>9 cells; lines = nuclear envelope). Cells were imaged at day-1 after FD-induction, when FD’s remain sparse. **i)** FD-to-nucleus distance fluctuates over time as an FD moves (n=4 cells; star = rupture). **ii)** GFP-KU80 kinetics shows rapid mislocalization after FD contact (∼min’s) followed by a slow recovery (∼hr’s). Red lines indicate duration of FD contact with the nucleus. **D**. FD-loaded cells exhibit ∼2.5-fold higher mislocalization of KU80 compared to control cells over the course of live imaging (n> 330 cells) or in fixed samples (n>500 cells; p<0.05), respectively.

To further assess the spatiotemporal dynamics of cGAS accumulation at a rupture site, we used a high curvature AFM tip (<100 nm diameter tip) to focally indent and rupture nuclei (**Fig.2B-i**). cGAS spreads again to a distance of several microns from the site of contact (**Fig.2B-ii**), and faster cGAS spreading might reflect a longer duration of rupture induced by the very high curvature AFM tip. Regardless, the cGAS pattern otherwise mimics focal rupture by small FD’s (**Fig.2B-iii, Fig.2A-i,ii**).

Nuclear exit of a diffusible DNA repair factor was our next focus in live cell imaging of FD-induced rupture, and GFP-KU80 was chosen because it is predominantly nuclear but can transiently mis-localize to cytoplasm upon nuclear rupture (**Fig.2C**) (Irianto et al., 2017), (Xia et al., 2018). Small FD’s within a few µm’s of the nucleus edge were tracked over time and showed modest changes in FD-to-nucleus distance over ∼10 min timescales, suggestive of oscillatory FD motility (**Fig.2C-i**) (Valm et al., 2017), (Welte, 2009) – although it is possible that immobilization occurs at later days when FD’s accumulate and jam together within the cytoskeleton (**Fig.1B-iv**). FD’s that transiently localized at the nuclear rim then often paralleled a sudden decrease (within ∼15 min) in GFP-KU80 nuclear intensity simultaneous with an increase in GFP-KU80 cytoplasmic intensity (**Fig.2C-ii**). Nuclear signal recovered after ∼1-2 h and remained constant, slightly longer than GFP-NLS recovery in other cell models (Robijns et al., 2016; Xia et al., 2018). Although only about half of all FD-nuclear close-contacts showed nuclear rupture (**Fig.2C-i** stars), overall ∼10% of cells loaded with FD’s showed real-time mis-localization of GFP-Ku80 — significantly exceeding rupture frequency of unloaded cells (**Fig.2D**), while also consistent with fixed cell results and with live-tracking of rupture with cGAS (**Fig.2A-iv**). Importantly, all KU80 mis-localization events in live imaging were associated with FD’s < 3 µm, consistent again with a relation to Lamin-B1 dilution (**Fig.1C**), whereas low curvature indentation caused by large FD’s were not sufficient to rupture nuclei and mis-localize KU80.

### Fat droplets increase DNA damage and cell cycle defects in 2D and after 3D migration

We next hypothesized that FD-loaded cells with nuclear rupture would exhibit more DNA damage, based on loss of key DNA repair factors for hours as exemplified by KU80 and on our previous observations of increased DNA damage associated with nuclear rupture after Lamin-A knockdown in standard 2D culture (Xia et al., 2018). Immunostaining for the DNA damage marker γH2AX shows FD-laden cells with cGAS+ nuclear rupture maximize DNA damage (**Fig.3A**). In the absence of such rupture, DNA damage remained at a basal level regardless of FD’s, indicating that FD’s do not increase DNA damage through other mechanisms. Nuclear rupture in the absence of FD’s tends to slightly elevated DNA damage, but the higher frequency of rupture with FD’s (**Fig.2D**) would be expected to maximize damage. DNA damage can trigger checkpoint(s) that impede cell cycle as we have recently shown with Lamin-A suppression in beating hearts from chick embryos and in standard 2D cultures of cardiomyocytes (Cho et al., 2019). FD-loaded cells here indeed show an increased fraction of cells in S-phase (**Fig.3B**), suggesting that FD’s can initiate *indentation* → *damage* → *cell cycle perturbation* (**Fig.3C**). Rescue of rupture-induced DNA damage and cell cycle delay is indeed achieved in part by overexpressing KU80, KU70 plus one more repair factor (Xia et al., 2019).

**Figure 3.**
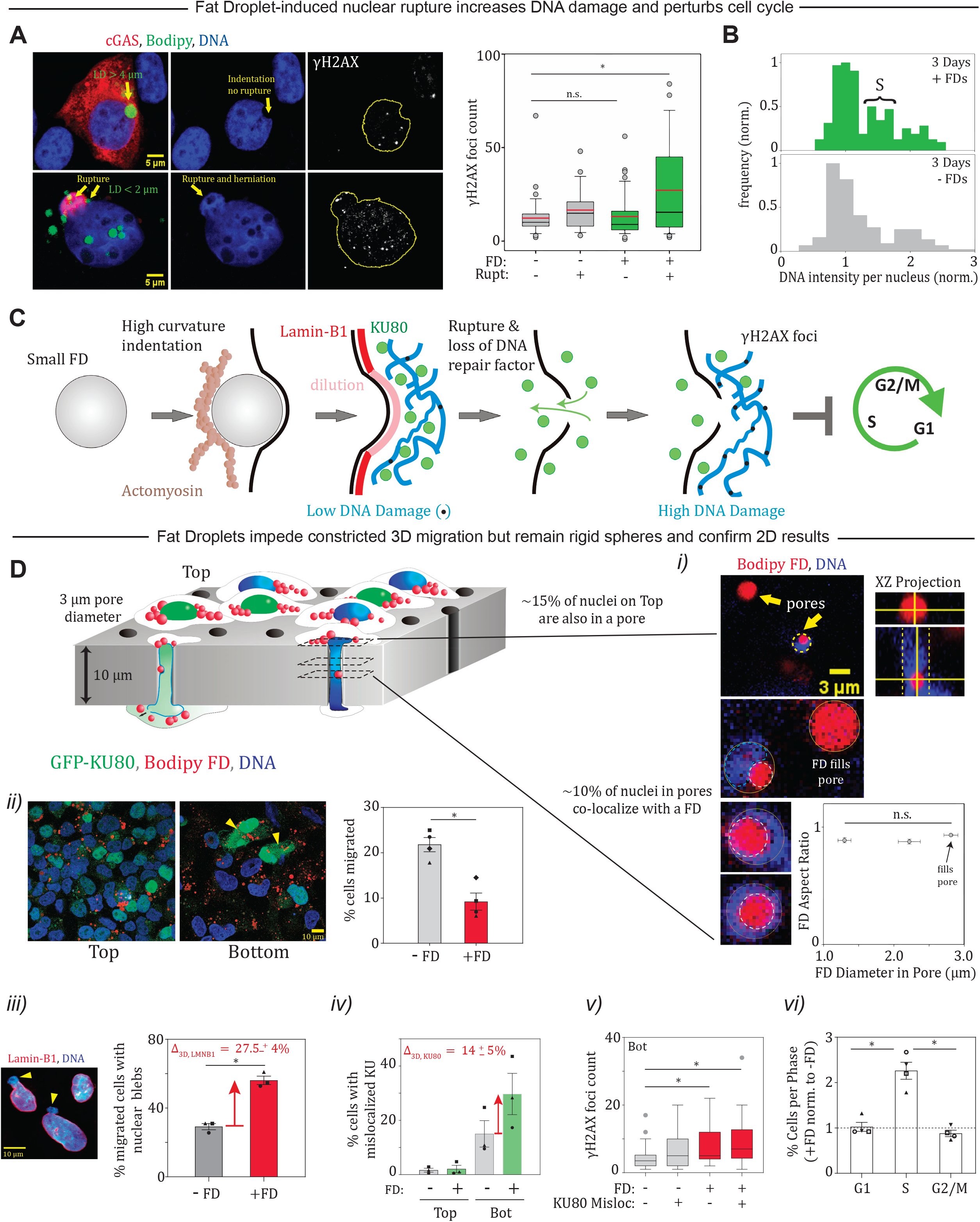
DNA damage and cell cycle perturbation are increased with FD-induced nuclear rupture, and 3D constricted migration confirms FD disruptive effects. **A-C**. Compared to non-ruptured controls, FD-ruptured nuclei (cGAS+) show more DNA damage based on IF for γH2AX foci 24 h after FD induction (n=138; p<0.05). Cell cycle of FD-loaded cultures is perturbed, with more cells in S phase after FD induction (n>350 cells). Doubling times for standard U2OS cultures is ∼1.5-days. *Pathway*: Small, stiff FD’s create high Gaussian curvature at indentation sites, which dilutes lamin-B1, favoring nuclear envelope rupture, rapid mislocalization of DNA repair factors, increased DNA damage, and perturbed cell cycle. **D**. Cells were densely plated on Top of Transwells at day-1 after FD-loading and allowed to migrate through 3D constricting pores (3 µm diam.) for 24 h more. **i)** Cross section confocal slices from within Transwell pores show FD’s remain round as they further distort the pore-deformed nuclei, which are pushed against the rigid pore wall (n=13) **ii-iv)** Fewer cells with FD’s migrate to Bottom versus controls, and yet ∼2-fold more FD-loaded cells show rupture with lamin-B1 deficient nuclear blebs (N=3 exp; n>600 cells; p< 0.05) and mislocalization of DNA repair factors (circle, square = KU80; triangle = KU70) (N=3 exp; n>360 cells). We must note that KU80 mis-localization between conditions on Top, with its high cell density, is less certain than on Bottom but seems lower in both than for migrated cells. **v, vi)** Migrated FD+ cells with nuclear rupture again show maximize yH2AX foci (n>120 cells; K-S test), and FD+ cultures again show more cells in S phase (3D = filled points, 2D = open; n>500 cells; p<0.05) with a tendency for G2/M suppression.

To further assess the effective stiffness of the small FD’s and their impact on other cell functions relevant to MSCs and some FD-containing cancers, such as migration (Cruz et al., 2020), we plated the U2OS cells with or without FD’s at high density on Transwell membranes with 3 μm diameter pores and then fixed, stained, and imaged by confocal microscopy after 24 h (**Fig.3D**). The vast majority of FD’s (>90%) are smaller than the chosen pore size (**Fig.1B-i**), and we find many examples of FD’s that co-localize with a nucleus within a pore (**Fig.3D-i**). Importantly, the nucleus deforms but the FD remains round, even for ∼2 μm FD’s that strongly distort the nucleus as the rigidity of the pore back-stops the nuclear indentation (instead of actomyosin stress fibers per **Fig.1B-iv**). The results further underscore the rigidity of FD’s.

Unlike very soft organelles (e.g. Golgi), rigid FD’s physically perturb actomyosin structures (**Fig.1B-i,iv**) which could impact actomyosin functions such as helping cells to pull themselves through constricting pores in the ∼3 hr’s it typically takes for migration from Top to Bottom of a Transwell (Xia et al., 2019). Fewer of the FD-loaded cells indeed migrated in the same period as control cells (**Fig.3D-ii**), with almost as much suppression as occurs with myosin-II inhibition of U2OS cells. Nuclear indentation by the rigid FD’s in pores might also impede migration, but the small fraction of such cells (∼10% of nuclei in pores; **Fig.3D-i**) would not account for the much larger effect on constricted migration.

Constricted migration through 3 μm pores is known to cause nuclear rupture, dysregulation of DNA damage & repair, and impede cell cycle after migration (Irianto et al., 2016),(Irianto et al., 2017), (Pfeifer et al., 2018), (Xia et al., 2019). FD’s increase *all* of these processes (**Fig.3D-iii,iv**), and ruptured nuclei again show maximum DNA damage (**Fig.3D-v**) with FD’s delaying cell cycle in the well-spread cells on Bottom per 2D culture (**Fig.3D-vi**). Generally, FD-induced scars of nuclear cGAS and the lamina are 2-3 fold more frequent than nuclei with KU80 mis-localization (**Fig.2D**,**Fig.3D-iii,iv**). Mis-localization of KU80 is less clear on the Tops, likely because cell density is so high (intentionally, to drive migration) that cells and nuclei remain rounded; this typically associates with limited acto-myosin stress fiber formation and would predict limited FD indentation (per **Fig.1B-iv**). Nonetheless, Lamin-B1 dilution can be found at sites of FD indentation, whereas nuclear wrinkles – which have no Gaussian curvature – show no Lamin-B1 dilution. The results overall provide further evidence for the indentation-damage-cell cycle checkpoint pathway caused by FD’s (**Fig.3C**).

### Interfacial tension is indeed high for FD’s in tissue adipocytes

Our various in vitro observations of small FD’s interacting with nuclei all indicate that small FD’s deform the nucleus upon contact rather than FD’s being distorted by the nucleus. This suggests that γ of FD’s in adipogenic cells is high and should be measured. Visceral fat tissue that was freshly isolated from mice was aspirated intact into micropipettes with supra-cellular diameters (∼50 - 100 μm) (**Fig.4A-i**) similar to those used to quantify the elasticity of embryonic heart and the viscoelasticity of brain (Cho et al., 2019). After a sufficient step in aspiration pressure, the tissue distends into the micropipette and then slowly flows; a creep compliance power law exponent α < 0.2 indicates a near solid-like response (**Fig.4A-ii**). Such isolated tissue has extracellular matrix (ECM) including multicellular fibers (**Fig.4A-i**) that likely modulate the mechanics (Heid et al., 2014) and that usually indicate a basal stress and a modest stiffness for a tissue (∼1 to 3 kPa) (Alkhouli et al., 2013); indeed, inverting the creep compliance values at 1 sec gives similar values and suggests tissue is overall stiffer than than the cells.

**Figure 4.**
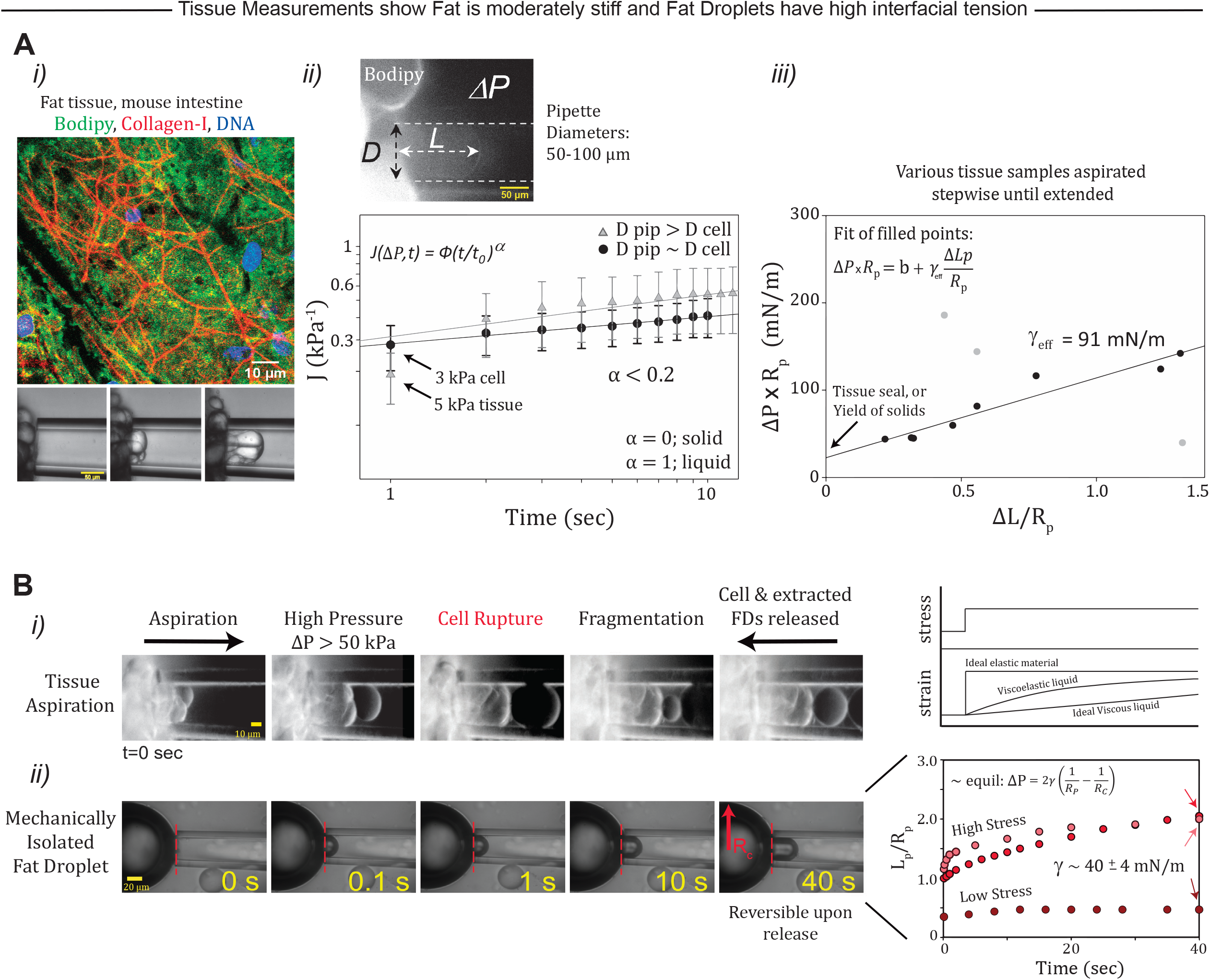
Fat tissue is solid-like and moderately stiff, based on micropipette aspiration, with mechanically isolated FD’s exhibiting a high interfacial tension similar to oil-water. **A. i)** Fat tissue resides within a collagenous network, as shown by a confocal image of fluorescently labeled mouse fat. During micropipette aspiration, individual fat cells are seen visibly entering the micropipette. **ii)** For analysis, tissue isolated from mouse intestines was pulled into micropipettes (diam. 50-100 μm) under controlled pressure *iJP*. The compliance of fat tissue exhibits a weak power law — a signature of a predominantly solid-like, viscoelastic response. **ii)** An effective interfacial tension γ_eff_ = 91mN/m was obtained from the linear fit of the tension versus strain measured across different fat tissue extracts (n=4 mice). The majority of the tissue measurements satisfy a linear response with R^2^ =0.93 (n=8; 3 outliers). **B**. Emulsions were formed from lipid content extracted from fat cells. **i)** Fat tissue was pulled into a pipette at high pressure (*iJP* > 50 kPa) causing membrane rupture and cell fragmentation before release. **ii)** Released lipids formed emulsions within the buffer-filled sample holder and the newly formed FD was subsequently re-aspirated into the pipette. *Top*: Under applied stress, a purely elastic material would instantly respond, and a viscous material would slowly respond. *Bottom:* Young-Laplace equation was applied to a near equilibrium strain, giving an interfacial tension of ∼40 ± 4 mN/m (n=3 FD’s).

The FD’s are nonetheless liquid based on molecular mobility measurements, and so if we assume that tissue stretching is resisted by an effective interfacial tension γ_eff_ (per spheroids, e.g. (Guevorkian et al., 2010)), then we find that single aspirated extensions of most tissue samples fit a line of slope γ_eff_ ∼ 90 N/m with an intercept that indicates a non-zero sealing pressure (or perhaps a yield stress) (**Fig.4A-iii**). The high value of γ_eff_ has complicated contributions from γ_FD_ and likely actomyosin as well as ECM. The softness of fat tissue (e.g. (Swift et al., 2013), (Alkhouli et al., 2013)), might be roughly estimated as *E* ∼ γ_eff_ / *R* ∼ 2 kPa based on *R* ∼ 50 µm for pipette size. For spheroids, the actomyosin term is ‘active’ in resisting high forces and increases from ∼5 mN/m at no stress to a high aspiration stress value of ∼25 mN/m (Guevorkian et al., 2010) – which is why we focused on low strain responses. Regardless, only a fraction of γ_eff_ should be conferred by that of the FD’s.

To directly measure γ_FD_ for fresh fat tissue – which is a measurement we were unable to find in the literature – we mechanically isolated large FD’s from adipose tissue via cycles of back-and-forth aspiration (**Fig.4B-i**). Once isolated, individual FD’s flow in aspiration as viscoelastic materials that nearly equilibrate in ∼10-100 sec (**Fig.4B-ii**). Release from aspiration shows fully reversible deformation, consistent with a γ that drives elastic recovery to a spherical shape of minimum area. Applying the Young-Laplace equation to distensions near equilibrium, allows us to measure a value of γ ∼ 40 ±4 mN/m (**Fig.4B-ii plot**). This is understandably lower than our γ_eff_ for intact fat tissue (**Fig.4A-iii**) and is higher than that of FD’s isolated from cultured Cos7 and HEK kidney cells and from fly hemocyte-like cells, but it is typical for oil-water interfaces (Israelachvili, 2011) (**Fig.1A-ii**). Our mechanically isolated FD’s from tissue may have distinct factors that affect γ, but – importantly – the result is consistent with FD’s having sufficient rigidity to remain spherical as they indent a nucleus.

## Conclusions

FD’s form in various cell types including some cancers and are viewed as potentially damaging (Cruz et al., 2020) – but a cell-type dependence could in particular reflect the ability of *small* FD’s to be suitably localized to a perinuclear location. Nuclear indentation by physiologically-relevant FD’s generates *inward* Gaussian curvature, as does AFM probing, whereas lamin-B1 dilution during nuclear rupture in other standard 2D cultures is observed at sites of *outward* curvature – and often with lamins that are mutated and/or suppressed (Xia et al., 2018) (Earle et al., 2020), (Pfeifer et al., 2022). The fit of our single filament model to the Lamin-B1 dilution results (**Fig.1C-iii**) is a first and anticipates rupture because similar parameters fit well to outward curvature rupture of progeria-derived MSCs in 2D culture (Cho et al., 2018) and of U2OS cells in transwell migration through pores (Pfeifer et al., 2022); the fit also suggests a complementary limit to lamin-A,C accumulation of several-fold, consistent with past results (Cho et al., 2018) but motivating further study. The lamina response to Gaussian curvature nonetheless begins to address a paradox as to why fat tissues, which are soft to the touch, possess high lamin levels – except in cases of lipodystrophy caused by lamin-A mutations or drug perturbations that keep it in a farnesylated form similar to lamin-B1’s. Evolution provides a useful perspective in that farnesylated Lamin-B1 is the ancient lamin that evolved to prelamin-A that is proteolytically cleaved to remove the farnesylation or else alternatively spliced as pre-mRNA to lamin-C that lacks a farnesylation site (Peter and Stick, 2012); in addition to this evolution via mutation and natural selection that overcomes a curvature-frustrating membrane interaction, regulation of lamin-A,C levels also evolved to be high in tissues when and where nuclear stresses are high (Swift et al., 2013). Such mechanoprotection is important because excess DNA damage can cause cell death (Roos and Kaina, 2013), but consistent with insights here, normal adipogenic differentiation shows that progressive *increases* in FD size (i.e. decrease in curvature) is accompanied by decreases in the levels and envelope localization of Lamin-A,C (Verstraeten et al., 2011). DNA damage modulates fat development (Nishio and Isobe, 2015), with adipogenesis involving withdrawal from cell cycle and DNA damage (Hatzmann et al., 2021); and DNA damage likewise enhances MSC osteogenesis – at least with nuclear rupture after constricted migration (Smith et al., 2018).

Actomyosin is displaced and suppressed by FD accumulation, which can feed forward to favor adipogenesis (Spiegelman and Farmer, 1982). High actomyosin levels and forces favor osteogenesis (Dupont et al., 2011) as well as nuclear rupture, but whether nuclear rupture by FD indentation depends on some low level of actomyosin as a backstop (**Fig.1**) requires further study – especially with attention to microenvironment. Indeed, extracellular matrix (ECM) that is stiff increases actomyosin stress (Cho et al., 2019), and yet fat has intermediate levels of fibrillar collagens (Swift et al., 2013), including at least one collagen associated with obesity (Pope et al., 2016) (Rodriguez et al., 2017). These structural ECM proteins might also mechanosense fat rigidity that ultimately has its basis in FD’s interfacial tension γ – which is not small like that of protein condensates that distort within a cell (Boddeker et al., 2022), but is instead rather large in requiring >100-fold more work to distort FD’s in a cell (**Fig.1A-ii**).

## Materials and Methods

### Cell Culture

U2OS cells were cultured in DMEM high glucose medium (Gibco) supplemented with 10% FBS (Sigma) and 1% penicillin/streptomycin (Gibco). For adipogenic differentiation, bone marrow derived MSCs were plated (5000 cells per mm^2^) on 6 well plates and induced with standard adipogenic media for 2 weeks. All cells were kept incubated at 37 °C with 5% CO_2_.

### Droplet Formulation and Fluorescence Staining

Tris HCL-Hydroxymethyl Aminomethane Hydrochloride- and Tris Base (both from Fisher Scientific) were dissolved in deionized water and combined with Fatty Acid-Free Bovine Serum Albumin (Sigma). The resulting solution was supplemented with Oleic Acid (Sigma-Aldrich), well-mixed, and purified using a 0.20 micrometer syringe filter (Fisher Scientific). The resulting stock solution was mixed with complete DMEM media to obtain a final concentration of 1 mM. Bodipy dye (FL-C12 and 558/568) was added at a 1:1000 dilution to the media-oleic acid mix prior to media replacement into pre-seeded well plates. Cells were incubated in the mixture for a duration of 24-72 hours.

### Confocal and Fluorescence Microscopy

Lower magnification epifluorescence and brightfield images were obtained from an Olympus IX71 microscope with a 40x/0.6-NA objective and a digital EMCCD camera (Cascade 512B; Photometrics). Confocal images were captured using a Leica TCS SP8 system with a 63x/1.4-NA oil immersion objective. When mounting was necessary, fixed samples were placed between 2 glass coverslips wet with 15 μL antifade mountant (Prolong Gold & Prolong Diamond, ThermoFisher) and allowed to set overnight. Experiments involving fixed, wet samples were performed, stained, immersed in PBS, and imaged within 35-mm coverglass-bottom dishes (MatTek).

### Theory: *Single-filament model*

As previously described (Xia et al., 2019), (Pfeifer et al., 2022), our parsimonious model considers a single stiff lamin-B1 filament with an *in situ* length *L*_fil_ and persistence length *𝓁*_p_ (Turgay et al., 2017). The filament is either attached to or detached from the nuclear membrane; thus the following partition function for the filament is obtained:

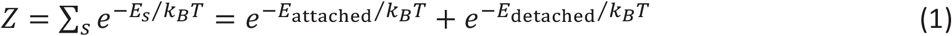

The detached state is taken as the reference state where *E*_detached_ = 0. For a flat nuclear membrane, *E*_attached_ equals the negative binding energy, -*E*, which favors filament attachment. However, since the nuclear membrane can be curved (with curvature = 1/*R*), two competing contributions to the energy of the attached state are considered:

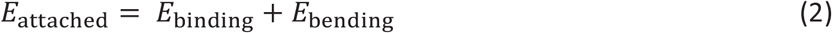

*E*_bending_, written as 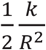 (where *k* is the filament bending modulus), accounts for the energy cost of bending the stiff filament along the curved membrane whereas *E*_binding_ comprises the negative binding energy which favors filament attachment. *E*_binding_ is modulated by membrane curvature because curvature can alter the filament-membrane contact area. Thus, Eq. 2 can be expanded to be written as a function of curvature:

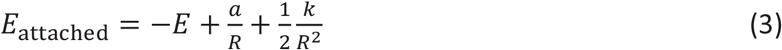

The *a*/*R* term, where *a* is constant, captures the change in contact area between the nuclear membrane and lamin-B1 filament resulting from induced curvature. The latter terms increase for decreasing R (i.e. higher membrane curvature), which means filament attachment becomes energetically costly at high membrane curvatures—and detachment more likely. The probability of the detached state is:

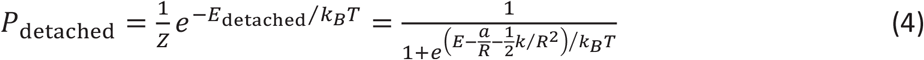

In the high-curvature limit, 1/*R* becomes very large and *P*_detached_ → 1, consistent with a high probability of lamin-B1 dilution and perhaps nuclear rupture at sites of FD-induced, high membrane curvature (Fig.1D; Fig 2A, C). In the low curvature limit, 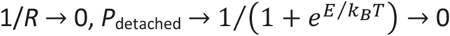, assuming *E* » *k*_*B*_*T*. A lamin-B1 filament is thus unlikely to detach or dilute when low curvature is introduced by a large FD.

In this model, the term *k* depends on filament length *L*_fil_ and persistence length *𝓁*_p_: *k* = (*𝓁*_p_*k*_*B*_*T*)*L*_fil_.The values *L*_fil_ = 0.38 µm and *l*_p_ = 0.5 µm from Turgay et al., 2017 where utilized to calculate *k* = (*l_p_ k_B_ T*)*L*_fil_ = (0.5 µm)(0.38 µm)*k_B_ T* = (0.19 µm^2^)*k_B_T*. Additionally, the bending term 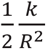 has almost no effect on the best-fit curve when fitting data, so we exclude it. Fits are thus of the form:

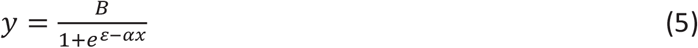

where *B* is a fit parameter and *ε* and α are a binding energy and an interaction energy, respectively. Here, FD’s that interact with the nuclear membrane impose a curvature *D*, where *D* is most commonly the diameter of the FD indenting the nucleus. Thus, the equation may be rewritten as:

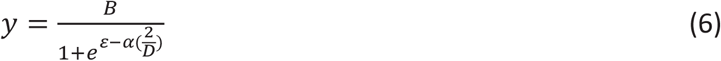

where curvature is defined as 1/radius. Thus, the size of FD indentation is correlated to lamin-B1 dilution. The measured dilution takes the form:

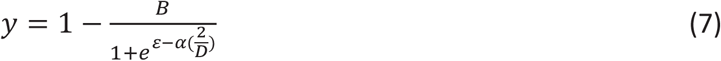

The fit to Fig.1C reflects: B=0.60, ε=4.8, α = 13.5. These parameters have very similar values to those applied to nuclear rupture (Xia et al., 2018; Pfeifer et al., 2022).

Observing that lamin-A responds inversely to lamin-B1, we use our single filament model to quantify the dilution ratio of lamin-B1 to lamin-A as:

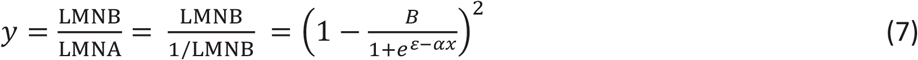

which is the square of the model. The first fit (blue curve) to Fig.1C inset is: B=0.50, ε = 4.8, α =13.3.

Finally, the model was used to predict the probability of lamin dilution via (5). The second fit (red curve) to Fig.1C inset uses: B=0.60, ε = 4.8, α =12.9.

### Live Imaging

Live imaging experiments were performed at 37 °C with 5% CO_2_ using an EVOS FL Auto 2 Imaging System with a 40x objective or a Zeiss Axio Observer 7 with a 40x objective over the course of 3-7 h. Plates were imaged every 5-10 minutes to minimize photobleaching and cell death.

### DNA Damage Assessment

Cells were fixed and immunostained with γH2AX. Foci were assessed and quantified in ImageJ using maximum signal intensity projections taken from raw confocal images stacks. Ten cells per field of view were randomly selected for manual counting of foci prior to application of a counting macro. The parameters of the counting macro which best reflected manual foci counts were applied to all other cell contours within the image.

### Transwell Migration

Constricted migration assays were performed using 24-well inserts with 3 μm diameter pores (Corning). 1.5 × 10^5^ cells were seeded on the tops of membranes in complete media. FD-loaded cells were plated on Tops of Transwells 24 hours after addition of oleic acid. Complete media was also added to the bottom of the insert such that there was no nutrient gradient. The assay was kept incubated at 37°C and 5% CO_2_ and allowed to proceed for 24 hours. After 24 hours, cells were fixed, and membranes were detached from the insert for immunostaining as described above.

### AFM indention during fluorescent imaging

Experiments were performed as described in Xia et al. 2018. Briefly, U2OS cells transfected with mCherry cGAS were replated on coverslips at a density of 60,000 cells/cm^2^ and cultured overnight. Coverslips were mounted in the fluid cell of a hybrid AFM system (MFP-3D-BIO; software: MFP-3D + Igor Pro 6.05; Asylum Research; Oxford Instruments) equipped with an inverted optical fluorescence microscope (Olympus IX81 with 40×/0.60 NA objective). Experiments were performed in a closed liquid cell at a temperature of ∼29°C in DMEM high-glucose medium with 10% serum buffered at pH 7.4 with 25 mM Hepes to prevent cell death in the absence of CO2. Cells were indented using MSCT-AUHW (Bruker) cantilevers with nominal spring constant 0.03 N/m, nominal tip radius 10–40 nm, and nominal tip height 2.5–8 μm. The cantilever spring constant was calibrated using thermal fluctuation method. The cantilever was positioned on the top of the nucleus, and the nucleus was indented with forces of ∼10–30 nN. When the cantilever deflection reached a predefined set point, the tip would dwell on the spot for 100 s before the cantilever was retracted and detached from the cell. Simultaneous fluorescent images were captured every 10 seconds for the entire probing cycle, including before the force was applied and after the cantilever was retracted for several minutes.

### Micropipette aspiration of fat tissue and mechanically isolated fat droplets

Fat tissue was excised from mouse intestine and immediately pulled using a homemade micropipette aspiration setup. Micropipettes were pulled from glass capillaries (World Precision Instruments, Sarasota, FL) with 1 mm inner diameters using a Flaming-Brown Micropipette Puller (Sutter Instrument, Novato, CA). Pulled tips were cut with ceramic tile to final inner diameter 50-100 µm. Pipettes were filled and treated for 20 min with 3% BSA in PBS to prevent sticking to the inside of the pipette. Pipettes were attached to a water-filled manometer-double reservoir of adjustable height. Pressures were applied with a combination of syringe suction and were measured using a pressure transducer (Validyne, Northridge, CA, USA) Imaging was done on a Nikon TE300 epifluorescence microscope with a 40×/0.60 NA air objective and a digital EMCCD camera (Cascade, Photometrics). Fat tissue was placed in a glass chamber filled with CO2 independent media and was aspirated at room temperature with pressures in the range of 1-50 kPa. The effective Young’s modulus of the aspirated spheroids was obtained from the linear relationship between the pressure and the strain *L*/*R*p for *L* ≤ *R*p,

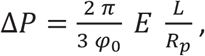

where ΔP is the pressure difference inside the pipette relative to outside, *L* is the length of tissue aspirated measured from the mouth of the pipette, Rp is the pipette’s inner radius, and *cp*_*o*_ is a shape factor ∼2 When a cell was aspirated at high pressure >50 kPa, cells rupture and the lipid is extracted and released into sample holder. The fat droplets that is formed in that way is aspirated again and the interfacial tension was calculated using Laplace’ law.

### Statistics

Analysis and model fitting was conducted using Prism (Graphpad) and SigmaPlot (SPSS). Statistical comparisons were conducted using two-tailed Student t-tests whereas one sample t-tests were conducted for normalized data. K-S tests were used to compare foci distributions. Tests were considered significant at P<0.05. Unless otherwise indicated, plots display mean ± SEM, with N indicating number of experiments and n indicating number of cells, objects, or measurements.

## Acknowledgements

The authors appreciate comments on the early figures and manuscript from multiple colleagues. Studies here of LD’s in adipogenic cells were subsequently followed up on by colleagues who studied hepatocytes (R.Wells et al). We thank the Penn Cell & Developmental Biology Microscopy Core’s Dr. Andrea Stout and Jasmine Zhao for microscope access and expert technical assistance. M.P.Tobin is supported by the National Science Foundation Graduate Research Fellowship Program under Grant No. DGE-1845298. The authors declare no competing financial interests. Grants from HFSP, NIH, NSF, HRFF, and others supported this work.

